# An Accurate Free Energy Method for Solvation of Organic Compounds and Binding to Proteins

**DOI:** 10.1101/2020.05.26.116459

**Authors:** Omer Tayfuroglu, Muslum Yildiz, Lee-Wright Pearson, Abdulkadir Kocak

**Author notes:** Corresponding to: Abdulkadir Kocak.

## Abstract

Here, we introduce a new strategy to estimate free energies using single end-state molecular dynamics simulation trajectories. The method is adopted from ANI-1ccx neural network potentials (Machine Learning) for the Atomic Simulation Environment (ASE) and predicts the single point energies at the accuracy of CCSD(T)/CBS level for the entire configurational space that is sampled by Molecular Dynamics (MD) simulations. Our preliminary results show that the method can be as accurate as Bennet-Acceptance-Ration (BAR) with much reduced computational cost. Not only does it enable to calculate solvation free energies of small organic compounds, but it is also possible to predict absolute and relative binding free energies in ligand-protein complex systems. Rapid calculation also enables to screen small organic molecules from databases as potent inhibitors to any drug targets.

## 1. Introduction

The novel coronavirus, SARS-CoV-2 (Severe Acute Respiratory Syndrome Coronavirus 2) is a causative agent of Coronavirus Disease 2019 (COVID-19).^1^ COVID-19 is now a global pandemic with unbearable economic burden and puts life of millions at risk around the world. The life cycle of the virus as many other viruses has an essential proteolytic auto-processing step.^2 3–4^ The 3CL-protease takes an important role in freeing individual functional proteins from single-chain polyprotein that has been translated by the host cell translation machinery.^5^ Interfering to this step has been shown to have capability of inhibiting virus replication.^6^ Therefore, 3CL-protease is one of the valid and most attractive drug targets for Covid-19 treatment.

Developing a drug molecule from scratch for curing a disease is quite expensive and time consuming. Instead, repurposing an FDA approved drug is a much faster and less expensive strategy since the time is very critical parameter in controlling such invasive viruses. Even testing all FDA approved drugs experimentally one by one may cause intolerable waste of the time. To help to ease the problem, computational calculations can be very handy. There have already been several computational studies about predicting an efficient FDA approved drugs that can be repurposed for COVID-19.^7–17^

In searching for drug candidates as inhibitors, there are two core questions that can be addressed: (a) what is the binding mode? (i.e. where does it bind from the enzyme?) (b) what is the binding affinity? (binding free energy).^18^ Inhibition of an enzyme can be involved in various mechanisms. For instance, an inhibitor can bind to the enzyme from the active site in the absence of substrate (competitive inhibition) forming an additional or it can bind in the presence of the substrate (i.e. enzyme-substrate complexed) from the allosteric site (uncompetitive inhibition). Alternatively, it can reduce the activity of the enzyme by binding from allosteric site of the enzyme either free or complex (non-competitive inhibition).^19^

In all these mechanisms the equilibrium constant, K_s_ belonging to the reaction between the enzyme and substrate is shifted towards the reactants by existing inhibitor in the media. Kinetic studies to understand the activity loss of the enzyme on substrate by the inhibitor report values such as inhibition equilibrium constant, K_i_ or half maximal inhibitory concentration, IC50. Both these experimental parameters are proportional (they are equal in special cases). Thermodynamic equilibrium constant K_i_ is related to the standard binding Gibbs free energy of the system by: ΔG^o^=-RTln(K_i_)

The ΔG^o^ can also be calculated from the thermodynamic potentials. Thus, it is possible to compare experimental ΔG^o^ determined from K_i_ and theoretical ΔG^o^ predicted by thermodynamic potentials using computational methods. However, logarithmic relation between the K_i_ and ΔG^o^ complicates the reliability of the calculations. An error of 3-4 kcal is acceptable for the most computations, but this makes thousands of times less efficient inhibitor. Therefore, most computational methods fail to estimate the experimental binding affinity trend (relative binding free energies) among the inhibitors. Reproducing the absolute standard free energies of binding is even more difficult for the computational methods. Moreover, a trade-off between the computational accuracy and speed must be made.

However, almost all of the studies screen databases listing the FDA approved drugs at the molecular docking level, which is mostly based on geometrical alignment of a ligand into a binding pocket, namely the active site of 3C-like protease (3CL^pro^). Despite the speed of docking approach, they are very coarse methods and almost never predict the correct experimental binding affinity trend among the inhibitors.^7–17^

More sophisticated methods to calculate the potential binding free energy of inhibitor candidate to the protein ranges from post molecular dynamics simulations such as Molecular Mechanics Poisson-Boltzmann Surface Area (MMPBSA)^20–23^ to perturbation methods such as Bennett acceptance ratio (BAR),^24–31^ the latter being much more accurate.

Although calculation of MMPBSA energies is quite fast as it requires only one MD simulation for each Protein-Ligand (PL) complex system and thus applicable to screen many drug candidates, it lacks of explicit water definition and uses the energies at the Molecular Mechanics (MM) level, very low level when compared to QM level definitions.

Perturbation methods like BAR requires several intermediate lambda (λ-) states in which the ligand can be decoupled, annihilated, or pulled. It uses the explicit water definition and its prediction for solvation free energies is quite successful. However, it still uses MM energy terms, and requires non-physical intermediate states. There have been several attempts to correct the MM energy in the BAR calculation by re-weighting the end-states with QM-MM definitions. In these post-BAR methods such as NBB, BAR+TP, only part of the system could be defined at the QM level.^26, 29, 32–36^ It could bring very little improvements in to the solvation free energies estimated by BAR.

In addition to BAR and post-BAR methods, others such as Free Energy Perturbation (FEP), Thermodynamic Integration (TI),^25–26, 31, 37–41^ Multistate BAR (MBAR)^42–44^ are also commonly used and proven to be sufficiently accurate in estimating solvation free energies along with relative and absolute binding free energies. However, all these methods require very high computational cost. Screening all the drug candidates using these methods may not be very practical. For a typical free energy of binding, these methods require at least 30 λ-states for the decoupling of L from PL complex and another that many for decoupling L from water (i.e. solvation free energy of ligand), which increases the computational cost by 60-folds.

Machine Learning (ML) techniques have attracted a great interest for the past decade due to their successful algorithms to scientific questions including but not limited to chemical reactions,^45–46^ potential energy surfaces,^47–50^ forces,^51–53^ atomization energies,^54–55^ and proteinligand complex scorings.^56^. One of the most promising aspects of the ML techniques is that the trained models can be applied to new systems (transferrable).^57^ Recently several models using active learning such as ANI-1,^58^ ANI-1x^59^ and ANI-1cxx^60^ have been trained to calculate DFT and CCSD(T) energies of small organic compounds containing C, H, O and N atoms in nonequilibrium conformations. It has been shown to reproduce the energies at the accuracy of the CCSD(T)/CBS level with billions of time faster than the actual QM calculations.^57^

Here, we introduce a new strategy to estimate free energies of solvation of small organic compounds and binding to proteins in explicit solvent using single end-state MD simulations. The method is adopted from ANI-1ccx neural network potentials (Machine Learning) for the Atomic Simulation Environment (ASE) and predicts the single point energies of the entire system at the accuracy of the CCSD(T)/CBS level. Our results show the method can be sufficiently accurate to measure solvation/binding free energies and fast making it applicable to screen FDA approved drugs.

## Theory

### Linear Response Approximation

The insertion of the ligand to an environment of solvent (solvation free energy) or receptor (binding free energy) can be defined by a coupling parameter, λ.^61^ At each λ-state, the potential energy of the system would be:

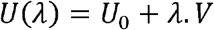

where V is the interaction energy between the ligand and environment. *U*_0_ is the reference potential energy, which includes all energy terms except for the ligand-environment interaction. The partition function is;

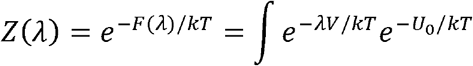

where F is the free energy depending on λ-state. Re-organizing the above equation,

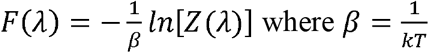

Thus,

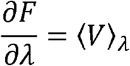

The equation above is only valid when the ligand-environment interaction is proportional to coupling term λ. This results in;

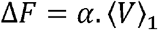

In the limiting case, this equation yields as;

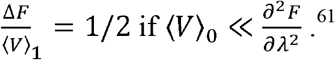

The equation implies that the free energy change is nearly half-the potential between the ligand-environment interaction in the complex configurational ensemble.^20^ So for the decoupling process;

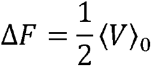

Here, we replaced the ML estimated CCSD(T)/CBS energies with MM energies at the λ=0 (i.e. coupled/bound) configurations. For each MD frame, three different energies were calculated ΔU_complex_, the ligand is complexed with environment (solvent or protein), ΔU_lig_, bare ligand that is extracted from the MD frame, ΔU_env_, the environment that is also extracted from the same MD frame which included the parts of either solvent (in the case of solvation free energy) or protein + solvent (in the case of binding free energy). And combining these three enrgy terms by ΔU= ΔU_complex_ – (ΔUiig +ΔU_env_). Finally, the averaged ensemble for the free energy was calculated by the standard methods introduced above using only one-endstate simulations. Interestingly, we found that experimental free energy (of solvation or binding) is approximately half of our calculated free energy. One should note that the energies calculated are of single point energies at the ML estimated CCSD(T)/CBS level in the clasical MD configurations. It does not include the Zero Point Vibrational Energy (ZPVE), entropy and PV terms.

## Methods

### System Preparation

For the binding studies, the crystal structures of 3CL^pro^ of SARS-CoV-2 of which the IC50 values are known were retrieved from Protein Data Bank (PDB). PDB IDs: 2GZ7, 4YOJ, 4MDS, 2ALV, 4WY3. Monomer structures were aligned and H-atoms were added using PyMol. The pKa values were calculated using Propka 3.1. The ligands were scanned for conformers. For the solvation free energies, we calculated molecules that have extensively available data in literature^34^ in order for a comparison. Using Gaussian 16 software, the ligands were first optimized at the B3LYP/6-31G* and ESP charges belonging to the conformer with the minimum energy for each ligand were generated at the HF/6-31G* level. Using Antechamber, RESP charges and GAFF force field atom types were generated. Acypype was used to convert Amber type input files to Gromacs.

### MD Simulations

The molecular dynamics simulations were carried out using Gromacs 2018 software package ^62^ with all-atom model of Amber ff99SB-ILDN force field ^63^ implemented in Gromacs.

For the binding studies, the protein-ligand complex (~4700 atoms) was placed in the center of a dodecahedron box. For the solvation studies, the ligand was placed in the center of a cubic box. Each system was solvated in the TIP3P model type water ^64^ with a cell margin distance of 10 Å for each dimension. The protein-ligand systems with ~70,000 atoms were neutralized and salted by 0.15 M NaCl. The neutral ligand only systems were not salted. Classical harmonic motions of all bonds were constrained to their equilibrium values with LINCS algorithm.

Energy minimization was carried out to a maximum 100 kJ.mol^−1^.nm^−1^ force using Verlet cutoff scheme. For both long range electrostatic and Van der Waals interactions, a cutoff length of 12 Å was used with the Particle Mesh Ewald method (PME) (6^th^ order interpolation).^65^ The neighbor list update frequency was set to 1 ps^−1^. As with our earlier studies,^66–67^ a stepwise energy minimization and equilibration schemes were used. Each minimization step consisted of a 5000 cycle of Steepest Descent and a subsequent 5000 cycles of l-bfgs integrators.

After minimization, each system was equilibrated within three steps using Langevin Dynamics. The first step consisted of a 1 ns of NVT ensemble. Experimental studies reported the solvation free energies at 298.15 K and binding free energies at 310 K. We have adjusted our simulations accordingly. The next steps consisted of NPT ensembles in which the systems were equilibrated to the 1 atm pressure by Berendsen for 200 ps and followed by Parrinello-Rahman isotropic pressure coupling for 1 ns to a reference pressure of 1 atm. When systems were reached to the equilibrium, an MD simulation of ~10 ns were carried out.

### Free Energy Calculations

For the BAR calculations, 10 equal decoupling steps (i.e. decoupling the ligand from the environment) of each coulombic and van der waals interactions, respectively (total 21 λ-windows) were used. For the BAR calculations of protein-ligand complex, a separate decoupling of the ligand from ligand+water system with the same parameters were performed and subtracted from the decoupling of ligand from protein+ligand+water complex system.^67^

MM-PBSA calculations were performed using g_mmpbsa script by Kumari et al.^21, 68^. For the calculations of protein-ligand complexes, we used the l=0 simulations of protein+ligand+water complexes. Only protein+ligand complex part from the trajectories were extracted and default parameters of implicit water with ε=80 and solute with ε=4 were used. Non-linear Poisson-Boltzmann equation with SASA model were used.

ML-QM calculations were performed using our custom build python scripts. For the calculations of solvation free energies, we used λ=0 simulations of ligand+water complexes whereas for the binding free energy calculations, we used λ=0 simulations of protein+ligand+water complexes.

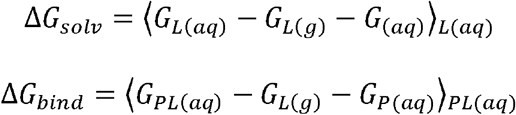

## Results and discussion

### Solvation Free Energies

In order to test the ML-QM method, we run 10-ns MD simulations for the molecules whose solvation free energies have been extensively studied. We first built a system in which the ligand is replaced in the center of box and the box is solvated with just single water molecule. The ligands studied are given in Figure 1.

**Figure 1.**
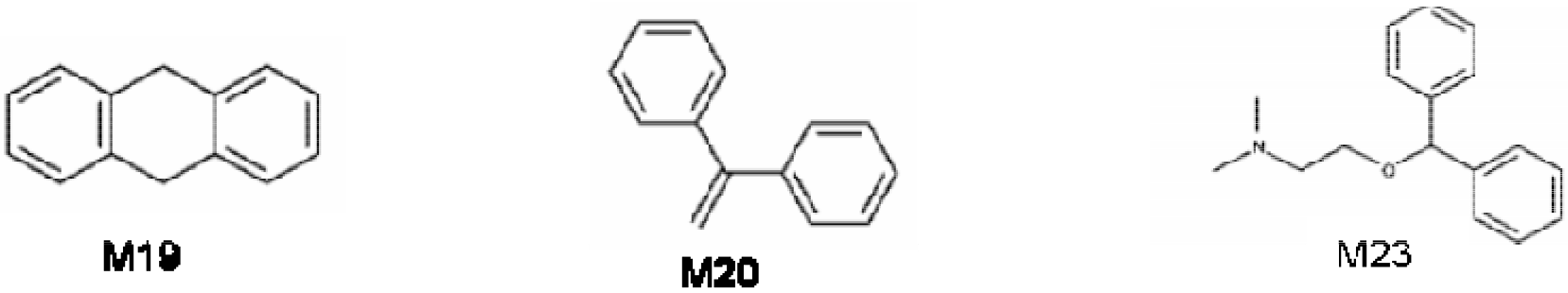
Molecules studied in this work for their solvation free energies

**Figure 2.**
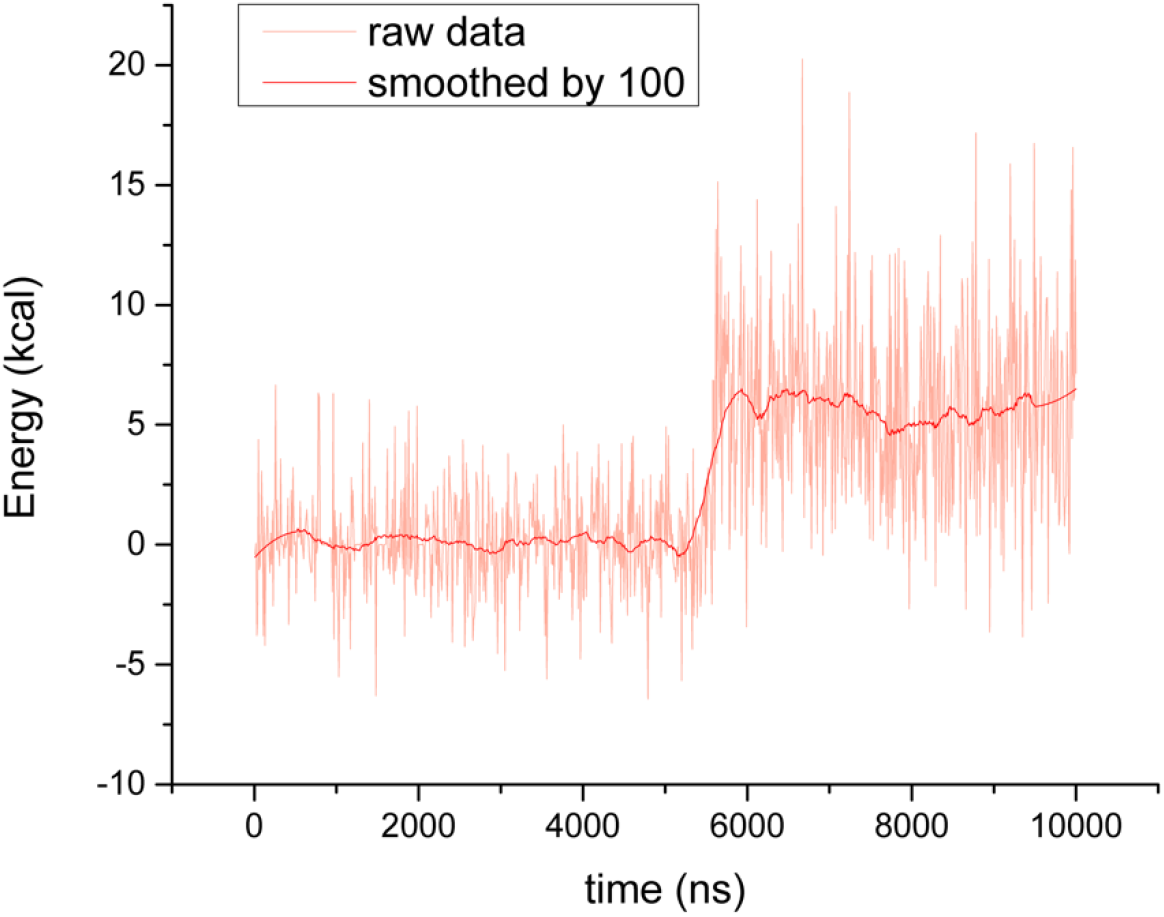
The ML-QM interaction energy calculated from the trajectory of MD simulation of M19 with 100 water molecules.

All the sampled configurations by the classical MD simulations were calculated for the single point energies using ML-QM method. Although the individual frames have very large energy fluctuations, the average energy of the system goes smoothly (Figure X)

Since the degree of freedom for the water molecule is much more in the case of unbound state in which water molecule is too far from the ligand and thus not interacting with the ligand, the MD could sample only unbound state. Therefore, ML-QM energy for the binding was almost 0. We prepared other independent simulations in which we increased the number water molecules (nH_2_O) in the box. Each time the average ML-QM energies were almost 0. This has been the case until nH_2_O=100. Figure 3a shows the interaction energy between the ligand and water molecules calculated by ML-QM energy difference: 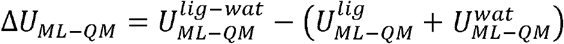 Here, 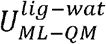 is the total energy of the system including ligand and water. 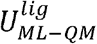 and 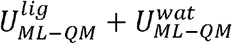 are the energies of free ligand and free water molecules, respectively, which have been calculated by extracting from the trajectory of complex structure. When we kept adding more water in the system and running other MD simulations, we found that starting from nH_2_O=100, the system undergoes from the unbound state to bound state by a drastic energy jump after ~9.5 ns. The degree of freedom for the water molecules in the bound state must be limited by the apparent interaction with the ligand. The same jump occurred for all of the other MD runs with increased amount of water molecules up to 200 (Figure 3b) but in the earlier time. Further addition of water molecules in the system causes the system to stay in the bound state all the time (Figure 3c). It is very apparent that the unbound states and bound states have the same averaged ensembles (Figure 3d).

**Figure 3.**
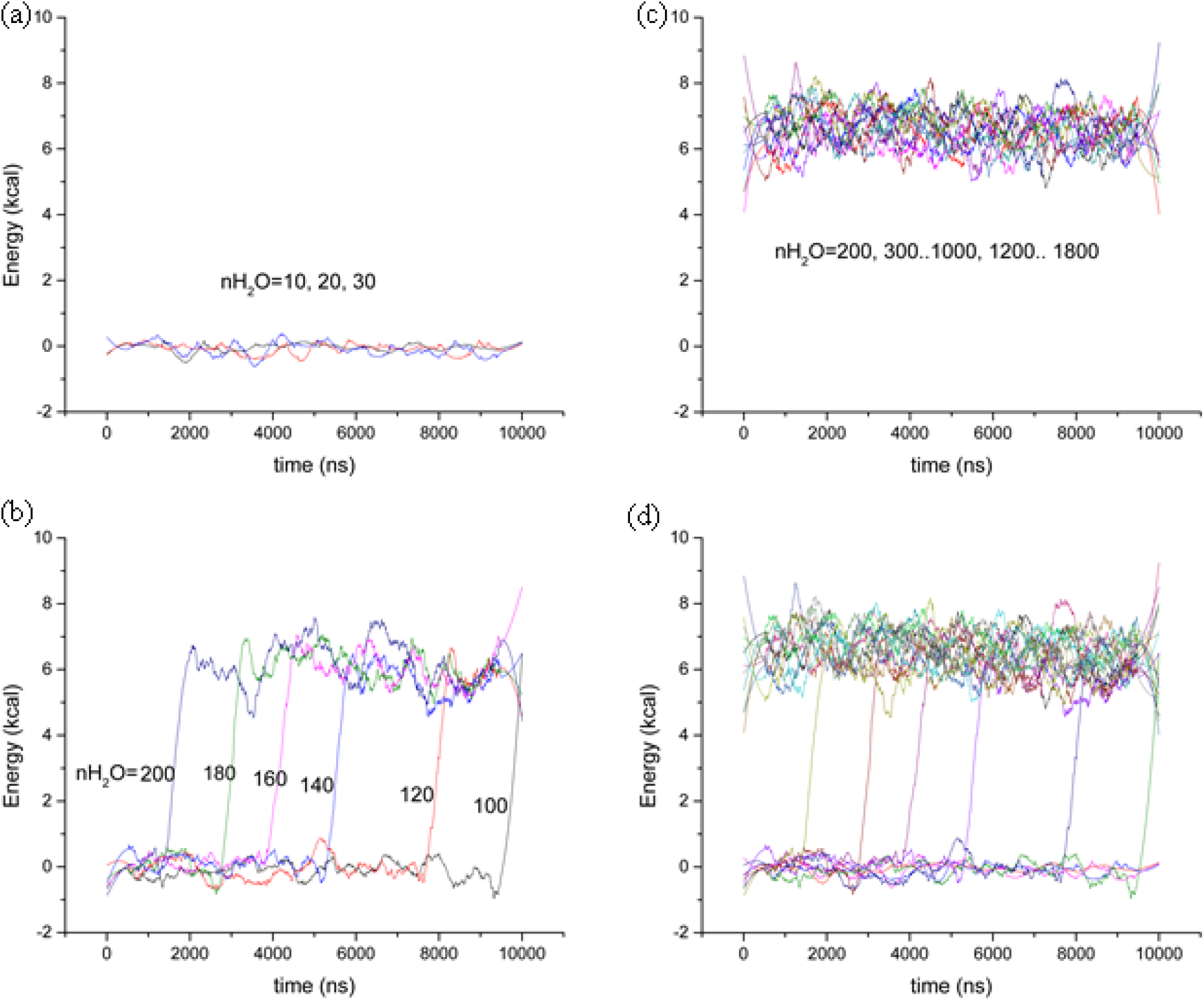
ML-QM interaction energy of M19 with different number of water molecules. Each data consists of the moving average of 100 points over 10,000 data set. (a) no interaction energy when the number of water molecules are much less than the configurational degree of freedom (b) the shift of the jump to an earlier time from unbound to bound state by addition of water molecules. (c) the system to stay in the bound state all the time when more than sufficient water molecules available. (d) the average energies of bound states to be all the same.

The average ML-QM energy of the entire MD simulation time can also be compared for each of the simulations with different number of water molecules. As in the case of simulations where bound-unbound state jump occurred, the energy profile of the average energies fit into a sigmoidal plot in which the inflection point corresponds to the experimental solvation free energies. The Figure 4 shows the average energies with respect to number of waters in each simulations.

**Figure 4.**
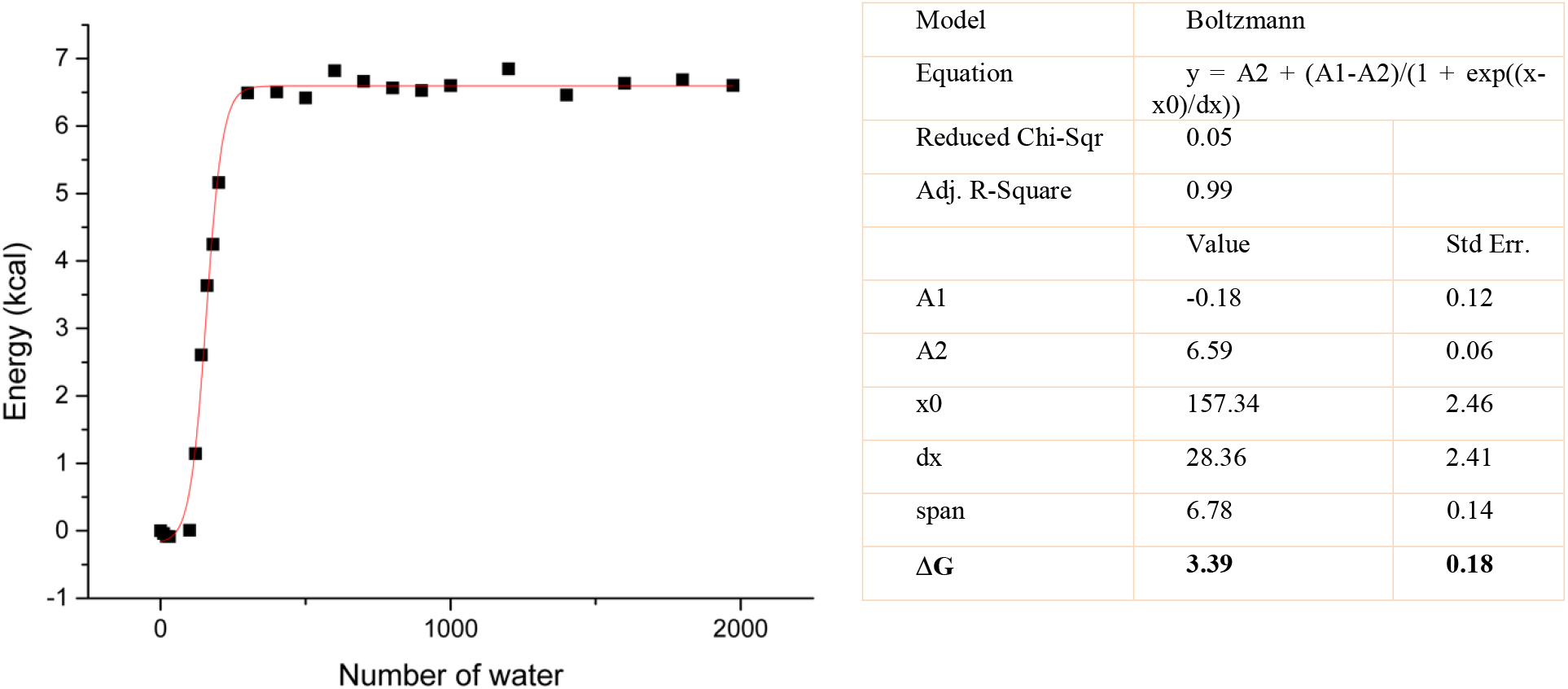
The ML-QM interacting energy depending on the number of water. The square marks show the average interaction energy over the entire MD simulations for each M19-nH_2_O complex (n=0, 10..., 30, 100, 120...200, 300...1000, 1200...1800, 1974)

It is apparent that the ML-QM interaction energy is constant for the simulations with more than sufficient waters exist. The empirically determined linearity constant is α=1/2, thus the free energy change is approximately half of the interaction energy between the ligand and environment.

Table 1 shows the experimental solvation free energies along with estimated by ML-QM and other techniques available in literature. ML-QM method significantly improve the results surpassing the conventional methods like BAR and MBAR.

**Table 1.** Experimental and calculated solvation free energies (kcal/mol). * values are from Ref.^34^

| Mol. ID | Experiment | ML-QM | BAR | BAR* | NBB* | BAR+TP* | MBAR* |
| --- | --- | --- | --- | --- | --- | --- | --- |
| 19 | -3.8±0.1 | -3.39 | -3.63 | -3.2 | -3.0 | -2.9 | -3.2 |
| 20 | -2.8±0.1 | -3.35 |  | -2.9 | -1.1 | -1.4 | -2.9 |
| 23 | -9.3±0.6 | -9.35 | -10.25 | -3.8 | -5.6 | -5.6 | -3.8 |

### Absolute Binding Free Energies

Using the limiting case, we could successfully calculate the binding of available inhibitors to 3CLprotease of SARS-CoV-2, of which the IC50 values available on literature. We compared the success of the method with other two common computational methods.

**Table 2.** Experimental inhibition profiles of the 3CL protease enzymes SARS-CoV-2 compared with absolute binding free energies. Experimental data were calculated by –RTln(IC50) at 310 K.

| PDB ID | Experimental | BAR | MMPBSA | ML-QM |
| --- | --- | --- | --- | --- |
| 2GZ7 | -9.25 | -9.78 | -39.88 |  |
| 4YOJ | -9.06 | -8.32 | -37.25 | -7.70 |
| 4MDS | -7.37 | -11.15 | -38.24 | -6.31 |
| 2ALV | -5.89 | -21.03 | -72.58 | -4.42 |
| 4WY3 | -5.13 | -9.82 | -56.63 | -9.37 |

It should be noted that the absolute binding free energy calculations may not directly reproduce experimental values, since experimental values are of IC50 values rather than the pKi values. However, considering the experimental conditions are more or less similar, the energy profile (i.e. the trend among the inhibitors) should at least be reproduced. Here, we see that ML-QM is the most accurate method to yield the correct trend among the compared methods (Figure 5).

**Figure 5.**
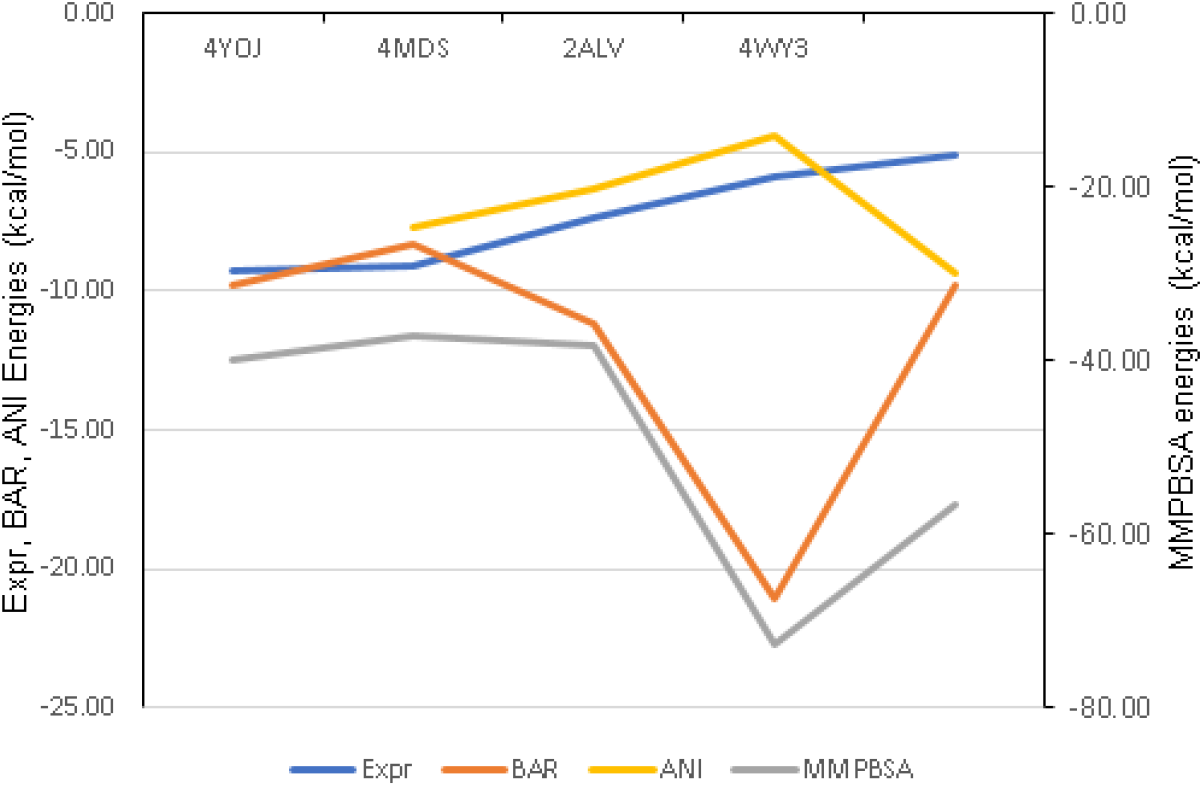
Calculated vs experimental binding profiles of inhibitors.

It should be noted that ML-QM and MMPBSA energies use single MD simulation trajectory, in which the ligand is fully interacting with the complex. The unbound states were created by removing the ligand from environment. Although the MMPBSA is as fast as ML-QM, the results are only meaningful in terms of relative binding energies. On the other hand, the BAR method uses 21 λ-states for the decoupling the ligand from the environment (protein and water) and another 21 λ -states for the solvation free energy of the ligand. The ML-QM surpasses both MMPBSA and BAR methods with increased speed and accuracy.

## Summary and Conclusions

Here we applied the Machine Learning techniques to calculate the experimental solvation and binding energies with increased speed at the accuracy of CCSD(T)/CBS level. The results show that the predicted binding energies are half of the interaction energies, which is a limiting case of LIE and reproduce the experiments better that well known conventional methods like BAR and MMPBSA. Since the method is fast and accurate, this method opens an area in the search of Covid-19 related inhibitors.

## Notes

### Competing Interest Statement

The authors have declared no competing interest.

